# Ecological specialisation and evolutionary reticulation in extant Hyaenidae

**DOI:** 10.1101/2020.10.14.338871

**Authors:** M V Westbury, Diana Le Duc, David A. Duchêne, Arunkumar Krishnan, Stefan Prost, Sereina Rutschmann, Jose H. Grau, Love Dalen, Alexandra Weyrich, Karin Norén, Lars Werdelin, Fredrik Dalerum, Torsten Schöneberg, Michael Hofreiter

## Abstract

During the Miocene, Hyaenidae was a highly diverse family of Carnivora that has since been severely reduced to four extant genera, each of which contains only a single species. These species include the bone-cracking spotted, striped, and brown hyenas, and the specialised insectivorous aardwolf. Previous genome studies have analysed the evolutionary histories of the spotted and brown hyenas, but little is known about the remaining two species. Moreover, the genomic underpinnings of scavenging and insectivory, defining traits of the extant species, remain elusive. To tackle these questions, we generated an aardwolf genome and analysed it together with those from the other three species. We provide new insights into the evolutionary relationships between the species, the genomic underpinnings of their scavenging and insectivorous lifestyles, and their respective genetic diversities and demographic histories. High levels of phylogenetic discordance within the family suggest gene flow between the aardwolf lineage and the ancestral brown/striped hyena lineage. Genes related to immunity and digestion in the bone-cracking hyenas and craniofacial development in the aardwolf showed the strongest signals of selection in their respective lineages, suggesting putative key adaptations to carrion or termite feeding. We also found a family-wide expansion in olfactory receptor genes suggesting that an acute sense of smell was a key early adaptation for the Hyaenidae family. Finally, we report very low levels of genetic diversity within the brown and striped hyenas despite no signs of inbreeding, which we putatively link to their similarly slow decline in N_e_over the last ∼2 million years. We found much higher levels of genetic diversity in both the spotted hyena and aardwolf and more stable population sizes through time. Taken together, these findings highlight how ecological specialisation can impact the evolutionary history, demographics, and adaptive genetic changes of a lineage.

## Introduction

Originating ∼25 million years ago (Ma) during the Late Oligocene ^1^, Hyaenidae was once a highly diverse family within the order Carnivora. During the Late Miocene there were dozens of species that occurred across Europe, Asia, Africa, and North America ^1^. However, this diversity has since been severely reduced to four extant genera, each of which contains only a single species, of which one ranges outside Africa (Fig. 1A). Extant hyenas consist of three species of bone-cracking hyena, the brown hyena (*Parahyaena brunnea*), the striped hyena (*Hyaena hyaena*), and the spotted hyena (*Crocuta crocuta*), and the insectivorous aardwolf (*Proteles cristata*). The aardwolf is unique amongst extant hyenas, not only in its feeding behaviour, but also in its morphology, and is believed to originate from “dog-like” hyena ancestors that were less derived than the extant bone-cracking species ^1^. Moreover, the oldest fossils that definitively show the specialized insectivorous traits of the aardwolf date back approximately 2 Ma, suggesting the aardwolf lineage only relatively recently started to evolve insectivorous adaptations ^2^. However, aardwolf-sized postcranial material dated to approximately 4 Ma suggests this process may have begun somewhat earlier. Taken together, these suggest that the current aardwolf morphology only evolved within the last 2–4 Ma and leads to the question of what underlying genetic mechanisms may have allowed the aardwolf to relatively rapidly evolve into a successful insectivore.

**Figure 1:**
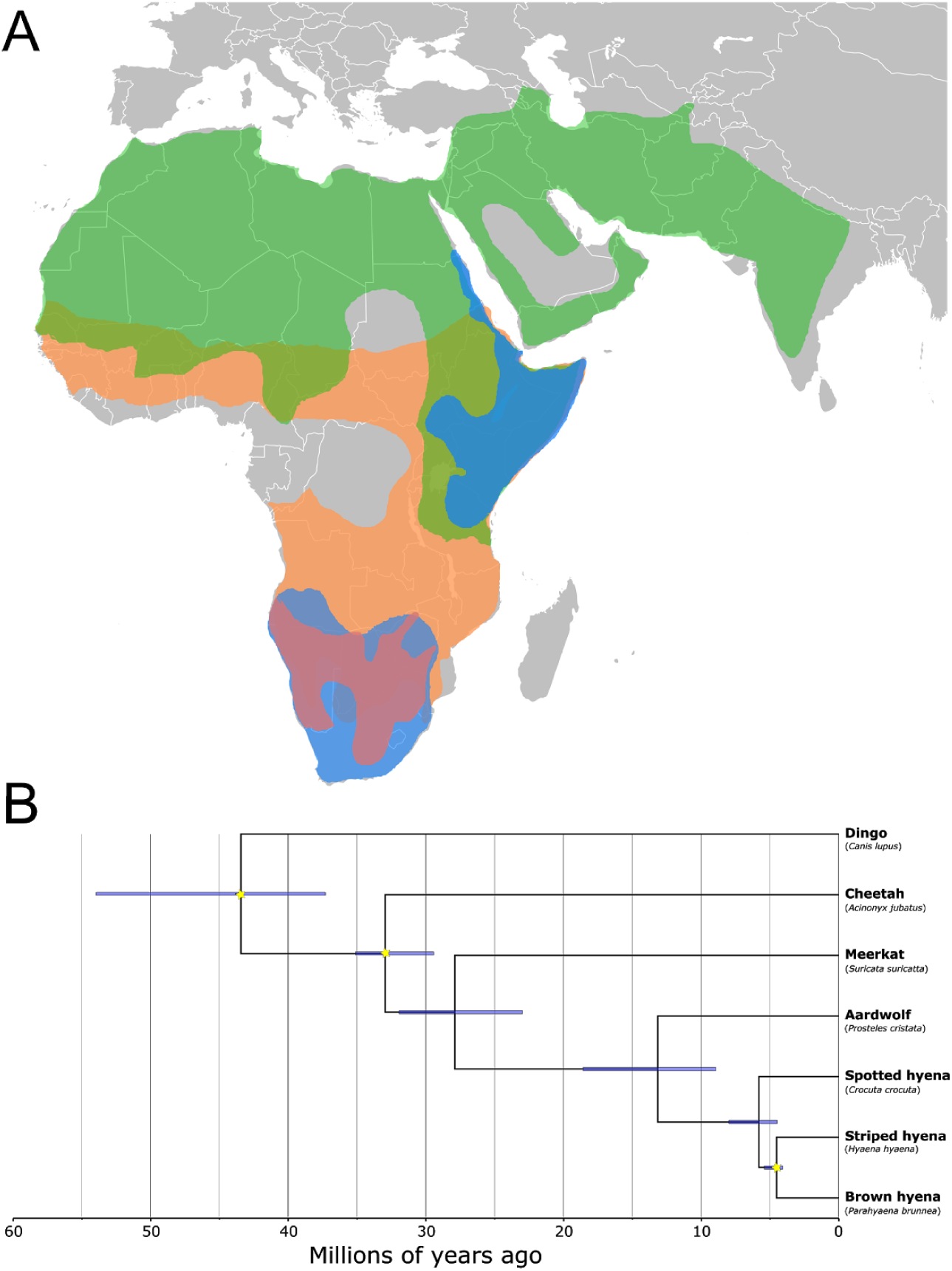
Distribution ranges and dated phylogeny of extant hyenas. **A)**Map showing the IUCN distribution ranges of the extant hyena species. Striped hyena − green, spotted hyena − orange, brown hyena − red, aardwolf − blue. **B)**Molecularly dated phylogenomic tree built using a Bayesian relaxed-clock analysis on the ASTRAL species tree using three fossil calibrations with soft maximum bounds (indicated as yellow stars). Blue horizontal bars represent 95% credibility intervals of node times.

The bone-cracking hyenas can further be separated into the more obligate scavenging species (brown and striped hyenas) and the spotted hyena, which largely relies on hunting but is also a facultative scavenger. These scavenging species play a vital role in maintaining the health of the ecosystem by facilitating nutrient cycling and influencing disease dynamics ^3,4^. However, despite their importance to their ecosystems, the genetic underpinnings of how these species adapted to such a potentially pathogen-rich feeding strategy remain unknown. Adaptations to this feeding behaviour may include highly sensitive olfactory senses ^3^and/or specific physiological mechanisms, like specialized digestive and enhanced immune systems ^5^. In addition to the uncertainty about the genomic underpinnings surrounding the adaptation of these species to their current lifestyles, their joint and respective evolutionary and demographic histories are still in question. Insights into these may help us understand how the extant species survived to the present day and also assess their resilience to future perturbations.

To this end, genomic analyses can provide profound insights into the evolutionary histories of extant species. To date, only two studies have investigated the evolutionary histories of species within the Hyaenidae based on genome data ^6,7^. One study on the brown hyena ^6^revealed exceptionally low levels of genetic diversity without any obvious detrimental effects on the survivability of the species, putatively caused by a long, slow and continual decline in effective population size (Ne) over the last million years. Another study, on the spotted hyena, revealed gene flow between modern African spotted hyena and Eurasian cave hyenas, an extinct sister lineage of spotted hyenas within the genus *Crocuta*^7^. These studies set the stage for future research on the evolutionary histories of extant Hyaenidae, including whether interspecies gene flow was a widespread phenomenon within Hyaenidae, and what role demographic history played in producing the diversity seen in these species today.

Here we set out to explore whole nuclear genomes to investigate the evolutionary histories of the remnant Hyaenidae species. We aimed to uncover whether interspecific gene flow was a widespread phenomenon within Hyaenidae, what genomic underpinnings may have allowed the aardwolf to become a specialised insectivore and the bone-cracking species to feed on carrion. Moreover, we set out to find the current levels of genetic diversity and offer explanations to current levels of diversity through investigating the respective demographic histories of each species over the last ∼2 million years.

## Results

### Evolutionary relationships

We inferred a dated phylogeny to gain insight into when the extant lineages within Hyaenidae diverged from one another (Fig. 1B). The aardwolf diverged first at ∼13.2 Ma (95% credibility interval (CI) 8.9 − 18.6 Ma), followed by the spotted hyena, ∼5.8 Ma (95% CI 4.5 − 8.0 Ma). The brown and striped hyenas diverged most recently ∼4.5 Ma (95% CI 4.1 − 5.4 Ma). However, such interspecific divergences are often complicated by post divergence gene flow ^8^. Therefore, to test for interspecific admixture within extant Hyaenidae, we implemented a three-taxon test for introgression known as D3 ^9^. We performed this analysis with every phylogenetically accurate triplet combination possible between the four hyena species (Supplementary tables S1 and S2). We found significant evidence for introgression between both the striped hyena/aardwolf (p=0.014 when mapping to the aardwolf genome and p=0.00022 when mapping to the spotted hyena genome) and the brown hyena/aardwolf (p=0.034 and p=0.00017) relative to the spotted hyena. We did not find evidence for different relative levels of admixture between striped hyena/aardwolf compared to brown hyena/aardwolf (p>0.05).

To further understand the evolutionary relationships among hyenas in light of this evidence of gene flow, we investigated potential incongruence among species tree estimates under different models of genome evolution. The phylogenetic estimate under the multispecies coalescent model placed the aardwolf as sister to the other three species of hyenas (Fig. 2A, i.). However, a locus concatenation method placed the spotted hyena as sister to the three other species (Fig. 2A, ii.). We further examined these results using the gene and site concordance factors of the data and inferences ^10^. Analyses of concordance factors showed that a majority of the whole gene alignments support the former resolution (48% versus ≤25% for other resolutions) (Fig. 2B). However, the majority of single nucleotide variants supported the latter resolution (45% versus ≤35% for other resolutions) (Fig. 2C). These results showed that a minority yet highly informative set of loci support the spotted hyena as sister to the three other hyenas.

**Figure 2:**
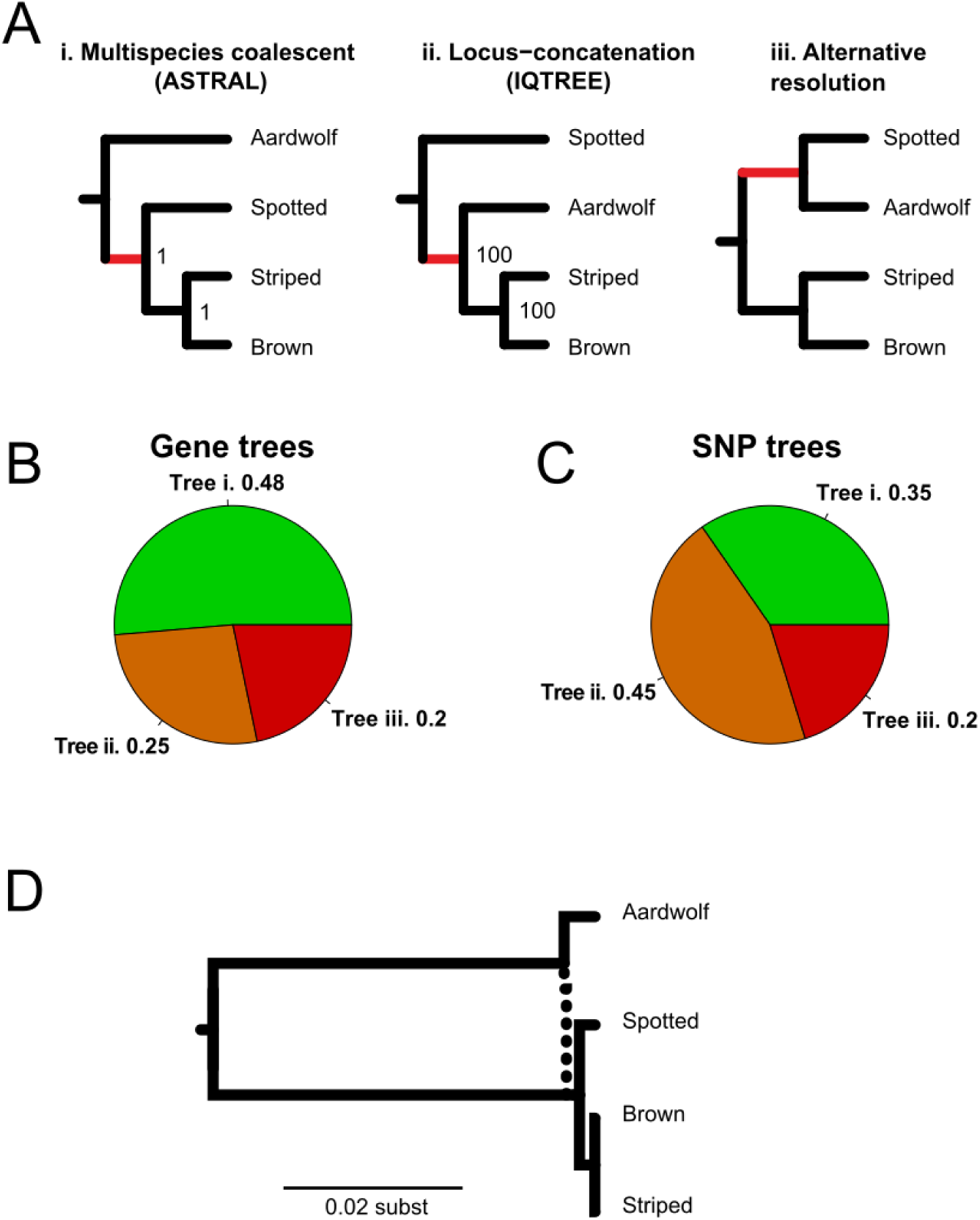
Support for the resolutions of the earliest divergence among extant species of hyenas. **A)**Fully bifurcating species tree estimates. Analyses assuming a fully bifurcating process support either the (i) aardwolf or (ii) spotted hyenas as the first to split from other extant lineages. Nodes in trees (i) and (ii) are labeled by their corresponding statistical support. **B)**Gene concordance factors for the three alternative topologies (i, ii, and iii). Numbers show the proportion of decisive gene trees containing that branch. **C)**Site concordance factors for the three alternative topologies (i, ii, and iii). Numbers show the proportion of decisive alignment sites supporting a branch in the reference tree. **D)**Multispecies network coalescent estimate calculated using PhyloNet allowing for a single introgression event.

Additional details on events of reticulation were explored via species tree reconstruction under the multispecies network coalescent. We used a Bayesian implementation of this model that samples gene trees with their coalescent times from sequence alignments ^11,12^. The analyses showed that the aardwolf originally diverged from other lineages of hyenas but underwent admixture with the ancestral lineages around the time of the split of the spotted hyena from the stem lineage of the bone-cracking hyenas (Fig. 2D).

### Genomic adaptations

To better understand which genetic adaptations occurred in Hyaenidae, we performed comparative genomic analyses to uncover genes under selection as well as a gene family expansion analysis on the olfactory receptors (OR). Out of the 9,400 1:1 orthologous genes in our data set, we uncovered 38 genes showing significant signs of positive selection in the bone-cracking lineage. In total, 35 of these had known annotations (Supplementary table S3). Furthermore, seven of these genes showed highly significant signs of positive selection (p >0.005). We also found signs of significant positive selection in 214 genes, 184 of which had known annotations (Supplementary table S4), in the aardwolf lineage. Of these, 60 were highly significant. Analyses of OR copy numbers revealed a substantial increase in both alpha and gamma family ORs in Hyaenidae (Supplementary figures S1 and S2). All hyenas showed at least 25–30% increase in alpha OR, and an approximately 45–50% increase in gamma OR compared to what can be seen in dogs. Furthermore, the alpha OR repertoire in hyenas was larger than observed in the other carnivoran species analysed here (cat, dog, and tiger). This result of a Hyaenidae-wide expansion in OR was confirmed using a subsequent *de novo*method implemented to avoid assembly or annotation biases (Supplementary figure S3). The same analysis also found expansions in immunity related genes in the bone-cracking lineage, and lipocalins and the UDP Glucuronosyltransferase Family (UGT) in the aardwolf lineage (Supplementary figure S3).

### Genetic diversity

We used autosomal heterozygosity as a proxy to estimate the genetic diversity of the four hyena species. We calculated autosomal heterozygosity using two different methods. The first method was implemented using ANGSD ^13^to allow for comparability to previous studies ^6,14^. The second method employed ROHan ^15^, which also calculates putative levels of inbreeding using runs of homozygosity (ROH). Based on the chosen method, absolute values were different but the relative numbers were comparable (Supplementary tables S5, S6, S7, and S8). The brown hyena had the lowest levels of heterozygosity, closely followed by the striped hyena, then the spotted hyena, and finally the aardwolf (Supplementary table S5). Relative to the other species included in this study, both the brown and striped hyena displayed very low levels, while the spotted hyena and aardwolf had medium to high levels of heterozygosity (Fig. 3A). We further analysed the distribution of heterozygosity across the genomes of the four hyena species in non-overlapping windows of 500kb (Fig. 3B). Both the aardwolf and spotted hyena had similar broad distributions across their genomes, although the spotted hyena had a much higher percentage of 500kb windows with less than 0.001% heterozygosity compared to the aardwolf. In contrast, both the striped hyena and brown hyena had much narrower distributions in heterozygosity with a skew in their distributions towards 0. The skew towards 0 led to further investigation whether inbreeding may have caused the low levels of heterozygosity within the brown and striped hyenas (Supplementary tables S6, S7, and S8). To explore this we calculated the percentage of the genome in ROH putatively due to inbreeding, specifying a 1Mb window with a heterozygosity proportion of less than either 1×10^−5^or 5×10^−5^. When specifying a value of 1×10^−5^, we did not find evidence for inbreeding in either the brown or striped hyena genomes. When specifying a value of 5×10^−5^, we did find some ROH in both the brown hyena (0.36% of the genome), and striped hyena (2.40%). However, both of these values were less than that found in the spotted hyena (5.12%). The aardwolf showed little to no signs of ROH regardless of the ROH heterozygosity value or mapping reference used.

**Figure 3:**
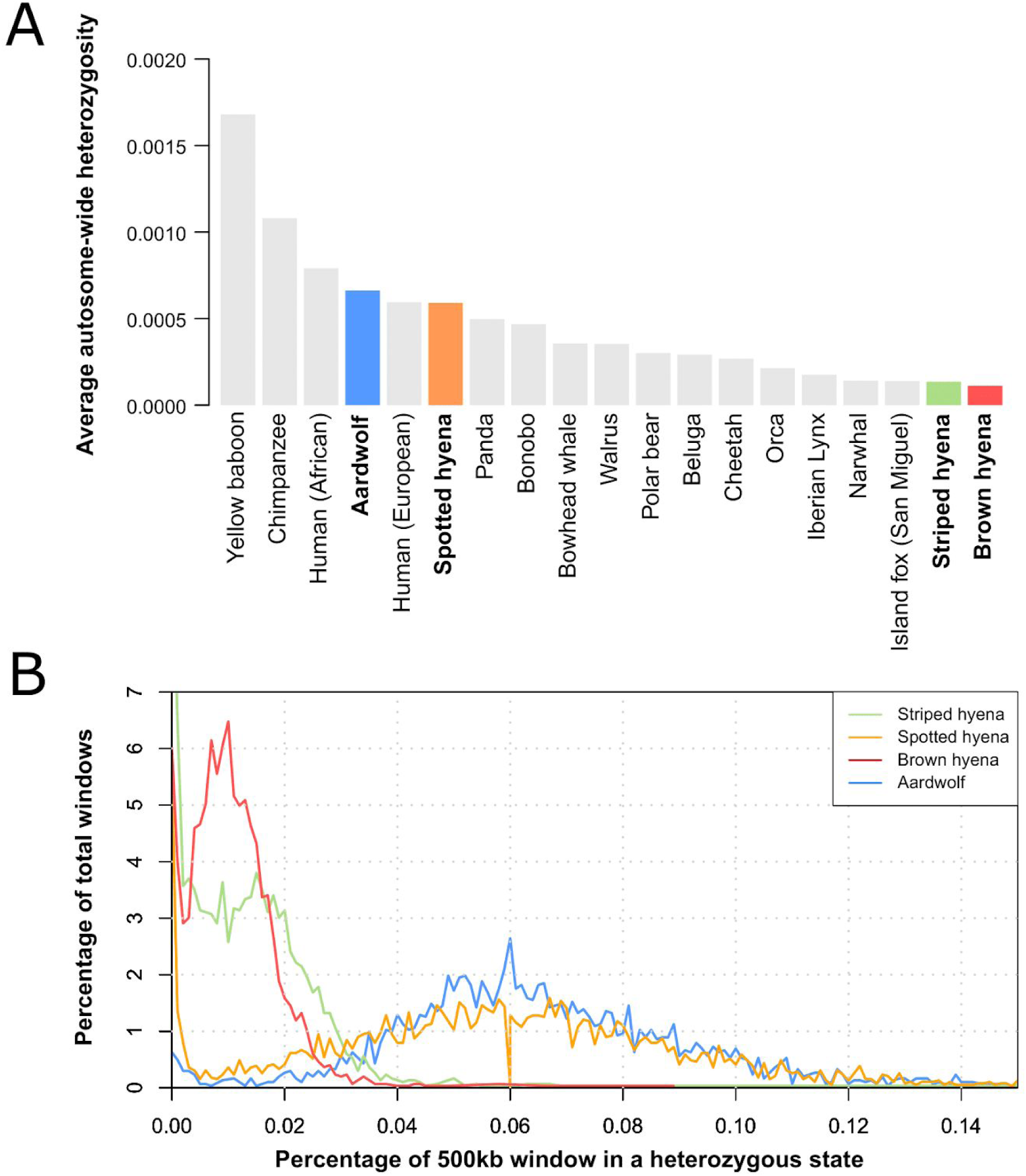
Comparative levels of heterozygosity within Hyaenidae. **A)**Average autosome-wide heterozygosity estimates for a number of mammalian species. **B)**Heterozygosity across the autosomes of the four Hyaenidae species calculated using 500-kb sliding windows.

### Demographic histories

To better contextualise the genetic diversity results we used a pairwise sequential markovian coalescence (PSMC) model ^16^and modeled the demographic history of each extant hyena species over the last ∼2 million years (Fig. 4). All species showed a unique pattern of demography over the last 2 million years but at the same time shared some similarities. The aardwolf showed the highest effective population size (Ne) through time relative to the other species with a relatively stable population size starting from ∼2 Ma until ∼500 thousand years ago (kya), where it began to increase until a sharp decrease at ∼100kya. The spotted hyena had the next highest Ne through time with a fluctuating Ne and a sharp decrease at ∼100kya. Both the brown and spotted hyenas show the lowest Ne through time. The striped hyena shows an increase in Ne at ∼500kya while the brown hyena continued to decline. Both showed a general, slow decline from ∼2 Ma, again followed by a sharp decrease ∼100kya.

**Figure 4:**
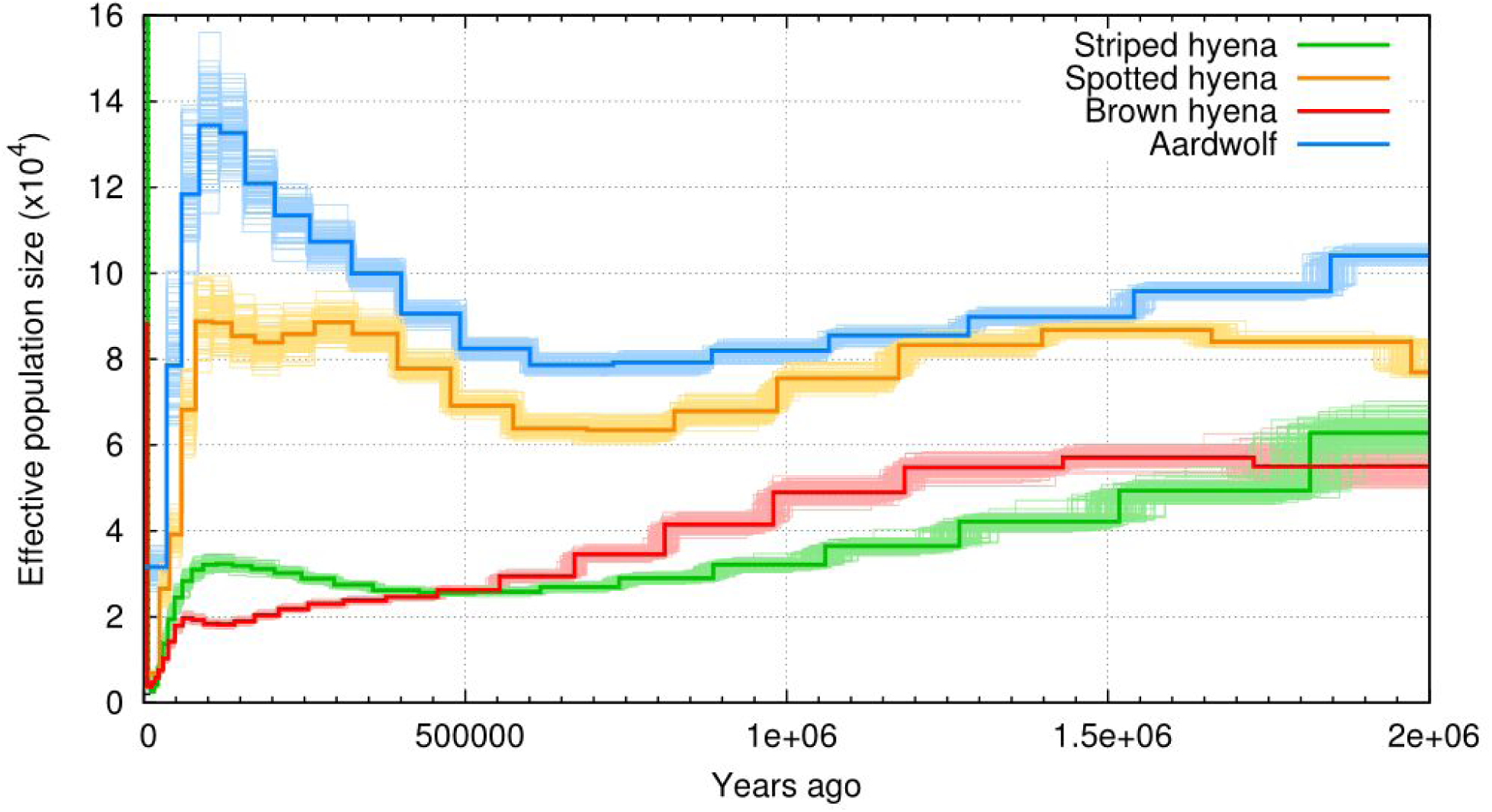
Demographic history of the four Hyaenidae species over the last 2 Myr, estimated using the Pairwise Sequentially Markovian Coalescent (PSMC) model.

## Discussion

Phylogenetic analyses revealed the same topology as that found in previous studies based on mitochondrial DNA and small numbers of nuclear genes ^7,17–19^. Our results confirm a comparatively deep divergence of the aardwolf from the other species (∼13Ma). This divergence date approximately coincides with the first appearance of Hyaenidae in Africa ^20^, tentatively suggesting that all living Hyaenidae today can be traced to an African origin. However, unlike previous studies we found a more recent divergence among the bone-cracking hyenas (∼6 Ma) (Fig. 1), albeit with overlapping credibility intervals. Our divergence date aligns much closer to the first appearances of canids in Africa ∼8 Ma (foxes) ^21^and *Eucyon*(the precursor to *Canis*) which appeared possibly as early as ∼6 Ma ago ^22^. The appearance of these competitors may have been the catalyst driving the evolution of the bone-cracking hyenas into the bone-cracking niche and allowed the persistence of these species while all dog-like hyenas (excluding the aardwolf) went extinct. Similar to the timing of the evolution of the bone-cracking hyena morph, our results suggest that the adaptation to insectivory of the aardwolf could also have been driven by competition from the expanding dog-like clades of canids. The resulting ecological specialisation may have allowed the aardwolf lineage to persist while all other dog-like hyenas were out competed. Both D3 ^9^and phylogenomic analyses supported an introgression event from the aardwolf lineage into lineages close to the divergence of the bone-cracking hyena lineages, and most likely disproportionately with populations of the stem ancestor of striped and brown hyenas. However, we note that this result may also arise due to introgression between the spotted hyena and an extinct unsampled sister lineage to the four hyenas studied here. This result is consistent with the relatively broad credibility interval of the timing of the split of the aardwolf from other lineages of hyena (Fig. 1), and the short branches in coalescent time units separating lineages of hyenas.

The deep divergence of the aardwolf and its substantially different ecology suggest introgression to be unlikely. However, considering our estimated divergence times between the lineages, especially with the 95% CI intervals of 8.9 Ma for the aardwolf (lower 5%) and 8 Ma for the spotted hyena (upper 5%), the divergence between the lineages when gene flow must have happened makes it not unrealistic. Moreover, there are no traces of any of the current aardwolf adaptations to insectivory prior to 4 Ma despite the extensive fossil record of Miocene hyenas. These observations suggest that an introgression event occurred ∼1.5M years after the spotted hyena diverged ∼6 Ma, but prior to the split of the brown and striped hyenas ∼4.5 Ma. This would also suggest that the specific insectivorous adaptations of the aardwolf did not start until the aardwolf lineage had ceased exchanging genetic material with the bone-cracking lineage.

The relatively rapid evolution of the aardwolf morphology within the last 2–4 Ma leads to the question of what underlying genetic mechanisms may have allowed the aardwolf to evolve into a successful insectivore subsisting on a diet almost exclusively consisting of termites ^23^. Investigations into the genes showing highly significant signs of positive selection in the aardwolf lineage revealed a number of genes putatively linked to aardwolf specific adaptations. We found five genes with known functions related to craniofacial development, putatively key in the formation of the unique aardwolf skull. These genes code for glycyl-tRNA synthetase (GARS), guanosine monophosphate reductase (GMPR), and stress-induced phosphoprotein 1 (STIP1), smoothened (SMO), and 3′-phosphoadenosine 5′-phosphosulfate synthetase 2 (PAPSS2). *GARS*loss-of-function gene variants have an impact on the developmental phenotype in humans including growth retardation, a large calvaria, and a high-arched palate ^24^. Less is known about *GMPR*, since fewer studies have focused on this gene. However, a 1 Mb *de novo*interstitial deletion in 6p22.3 chromosomal region, which contains the *GMPR*gene, was shown to cause some craniofacial malformations, suggesting there may be a link between this gene and craniofacial development ^25^. A large genomic screen identified *STIP1*to be in gene sets responsible for the branchial arch formation, also suggesting a role of this gene in craniofacial development ^26^. Conditionally null *SMO*mice had severe malformations in craniofacial structures developing from neural crest-derived elements ^27^. Mice having two-copies of an autosomal recessive mutant allele of *Papss2*have been characterized as having foreshortened limbs, a short stout tail, and a complex craniofacial phenotype ^28^. Furthermore, we also note one gene involved in skin barrier function (*DSC1*) ^29^. Skin barrier function may have been a key adaptation allowing aardwolves to transition to the insectivorous niche as the defense secretions of soldier *Trinervitermes*termites are known to be highly toxic, and despite this, the majority of the aardwolf’s diet stills consists of *Trinervitermes*^30^. Finally, evidence for an expansion in the lipocalin and the UDP Glucuronosyltransferase (UGT) gene families suggests important roles in the evolution of the aardwolf. Lipocalins produce major allergens that can induce severe anaphylactic reactions in humans when bitten by hematophagous insects ^31^while pseudogenisation of UGTs in the domestic cat are linked to an inability to process drug toxins ^32^. Although currently speculative, the aardwolf may have recruited these gene families as a defence mechanism against the termite toxins, enabling it to feed despite being bitten.

Our results suggest that the ability of the three scavenging hyena species to feed on potentially infectious carrion was facilitated by genetic adaptations to the immune system and digestion. A heightened immunity would allow these species to feed on rotten flesh containing many potential pathogens ensuring a relatively reliable food source ^5^. When fresh food was scarce, individuals that could survive on what was available may have had a competitive advantage over others. As immune-relevant genes are commonly found to be under positive selection due to their generally rapid evolutionary rate ^33^, it is difficult to discern whether these genes were flagged because of their faster background evolutionary rates or whether they are truly linked to adaptation to scavenging. However, our findings of a number of immune related gene families being expanded in the bone-cracking hyenas does add weight to the hypothesis that selection on immunity was key in the adaptation to scavenging. Despite these results, investigations into genes showing the most significant signs of positive selection, suggests that genes involved in the gastrointestinal system were under stronger selection than immune genes in general; in particular *ASH1L, PTPN5, PKP3*, and *AQP10*. In mice, *ASH1L*has been shown to be significantly downregulated in the colon during T cell-mediated colitis ^34^, and also has a potential connection to T cell-mediated autoimmune diseases. The *PTPN5*gene has been linked to viral gastritis in humans, an infectious disease of the gastrointestinal system that involves inflammation of the stomach lining caused by viruses (Genecards.org). Studies have shown *PKP3*to undergo molecular processes associated with inflammatory responses in the gut ^35^. Finally, *AQP10*appears to contribute to liquid transport in the gastrointestinal tract ^36^. All four of these genes suggest that specific gastrointestinal adaptations were vital in allowing hyenas to adapt to feeding on carrion. Moreover, in addition to their functions in the gut, variants of *PTPN5*have been shown to also cause increased bone mineralisation in mice (https://www.mousephenotype.org/) and therefore may have also played a role in strong tooth/jaw development required for bone cracking. Finally, *PKP3*has also been shown to also play a role in inflammation of the skin ^37^, an adaptation that may have also been vital to avoid disease while feeding on carrion.

The most parsimonious explanation for the observed expanded olfactory receptor (OR) repertoire in all extant species would be that it was also expanded in their common ancestor, and subsequently that an acute sense of smell may have been a driving force allowing for the survival of the family since the Miocene. This result was expected in the bone-cracking species due to their carrion feeding lifestyle as it could also be assumed that to find carrion before competitors, these hyenas would have required an acute sense of smell. However, the result was somewhat unexpected in the aardwolf as, on the surface, it could be argued that the aardwolf does not need as acute a sense of smell as the other species. However, the retention of this expanded OR repertoire may have been put to use in other ways (e.g. territory marking and mating ^38,39^), or it could have been maintained if there were no strong selective pressures against it.

While both the aardwolf and spotted hyena showed relatively high levels of diversity compared to a number of other mammalian species, both the striped and brown hyenas seem to have exceptionally low levels of genetic diversity. Similar to the brown hyena ^6^, this low level of diversity in the striped hyena was likely not caused by recent inbreeding, inferred from the low occurrence of ROH (Supplementary tables S6, S7, S8). These results raise the question which demographic changes through time led to the different present day genetic diversity levels in these species. Aardwolf and spotted hyena not only show relatively high levels of Ne through time, surprisingly, in light of their different ecologies, they also display similar fluctuations in population size, characterized by a slow decrease from ∼1.5 Ma followed by a recovery/increase ∼500kya. The striped hyena on the other hand shows a gradual decrease in Ne from about 2 Ma until plateauing at a low level ∼500kya. This is similar to what is seen in the brown hyena and may explain the ability of the species to retain such low levels of diversity without a detectable signal of inbreeding. We suggest that the parallel trajectories of genetic diversity in the striped and brown hyenas may be related to their similar social and ecological characteristics. Partly relying on carrion as well as being mostly solitary ^40^may have resulted in low carrying capacity and subsequently limited population sizes.

The striped hyena has the largest geographic distribution of all hyena species, ranging from eastern Africa to India ^41^. Very little is known about the range wide population structure and population densities of this species, but our results suggest that they may persist at very low densities across their range. Unfortunately, the genome used in this study was obtained from a sample of a captive individual (Tierpark Berlin.), wherefore it is unknown whether the genetic diversity of this animal reflects that of a wild individual. It was a minimum fourth generation zoo individual, with ancestors from Yerewan zoo, Bratislava zoo, and an unknown location. The potential mixed ancestry of this individual may provide a reason as to why there are no signs of inbreeding but if that was the case, the low heterozygosity found in this individual would be even more astonishing, with heterozygosity levels potentially even lower in wild bred individuals.

While the demographic trajectories between the high and low diversity species pairs were largely different through time, all four species showed a dramatic decline in Ne ∼100kya. A similar decline was also reported in a large dataset of African ruminant genomes and was proposed to have coincided with the expansion of humans and the resultant hunting of these large herbivores ^42^. Moreover, this was previously reported for the spotted hyena and was hypothesised to be linked to increased competition with humans and exacerbated by a decline in prey availability ^7^, which could also hold true for brown and striped hyenas. However, we also see a dramatic decline at approximately the same time in the aardwolf, a specialised insectivore expected to be in no direct competition with early humans ∼100kya. Even though humans may have hunted aardwolves, a more probable scenario would be that there were other environmental mechanisms besides humans (e.g. climate induced environmental change) leading to large decreases in Ne of African species ∼100kya.

## Methods

### Data acquisition

#### Aardwolf data generation

To generate a mapped aardwolf genome, we extracted DNA from a single aardwolf. The individual was captured for an ecological research project on Benfontein game reserve outside of Kimberley, central South Africa (28°52′ S, 24°50′ E) ^38^. The reserve covers approximately 11,400 ha and consists mainly of dry Karoo shrubland, arid grasslands, and Kalahari thornveld ^43^. The animal was initially captured in 2008 under a permit from the animal care and use committee of the University of Pretoria (EC031-07) and permits from the provincial government in the Northern Cape (FAUNA 846/2009, FAUNA 847/2009). Genetic samples consisted of a small piece of skin collected at the ear tip which was stored in 95% ethanol at −20°C.

We extracted DNA from the tissue sample using a Zymo genomic DNA clean and concentrator extraction kit (Zymo Research, USA), following the manufacturer’s protocol. DNA extracts were built into Nextera Illumina sequencing libraries (Inqaba Biotec, Pretoria) and sequenced on an Illumina Hiseq X at the NGI Stockholm, Sweden using 2×150bp paired-end reads.

### Previously published data

For use in our analyses we downloaded the previously published nuclear genome assemblies for the striped hyena (Genbank accession: GCA_003009895.1)^6^and the spotted hyena (Genbank accession: GCA_008692635.1) ^44^. We further downloaded the respective Illumina raw reads from each assembly as well as the raw reads from the resequenced brown hyena (Genbank accession: SRS2398897), which was previously studied by mapping to the striped hyena ^6^.

### Mapping of raw Illumina reads

Raw reads were all treated comparably before being mapped to a specific reference genome, which varies depending on the analysis. We used Cutadapt v1.8.1 ^45^to trim Illumina adapter sequences from the ends of reads and remove reads shorter than 30bp. We merged overlapping read pairs using FLASH v1.2.1 ^46^and default parameters. We mapped the trimmed merged and unmerged reads to the respective reference sequences using BWAv0.7.15 and the mem algorithm ^47^. We further processed the mapped reads using SAMtools v1.3.1 ^48^to remove duplicates and reads of low mapping quality (<30).

### Consensus sequence construction

For the comparative genome analyses, we used the two already assembled genomes (striped hyena and spotted hyena) and the two resequenced genomes mapped to the striped hyena (brown hyena and aardwolf). To decide which resultant mapping file to use, we mapped the brown hyena and aardwolf to both the striped hyena and spotted hyena assemblies and compared mapping statistics. As both species mapped more efficiently to the striped hyena genome (aardwolf − 457,078,015 reads / 72,698,723,765 bp mapped to striped hyena vs. 454,576,330 reads / 72,003,685,729 bp mapped to spotted hyena, brown hyena − 635,096,073 reads / 98,210,957,289 bp mapping to striped hyena vs. 613,078,338 reads / 94,209,031,605 bp mapped to spotted hyena), we built consensus sequences of these for further comparative and phylogenomic analyses. To build the consensus sequences we used a consensus base call approach (-doFasta 2) in ANGSD v0.913^13^and specified the following parameters: minimum mapping and base quality of 25 (-minQ 25 −minMapQ 25), only consider reads mapping to a single location (-uniqueOnly 1), and remove low quality secondary alignments (-remove_bads 1).

### Spotted hyena transcriptome

For the purpose of improving downstream whole genome annotations, we also included the assembled transcriptome of a single spotted hyena individual. Four tissues (brain, olfactory tissue and testis) of a male juvenile spotted hyena were sampled post-mortem, cooled, and frozen. From total RNA samples poly(A)-RNA was isolated, converted into cDNA by random-priming and cDNA libraries prepared using Illumina TrueSeq adapters by vertis Biotechnologie AG (Freising, Germany). Libraries were paired-end sequenced (PE 2×100bp and PE 2×150bp) at the Friedrich Löffler Institute (Jena) on the Illumina Genome Analyzer IIx.

We trimmed adapter sequences from the raw Illumina reads using Cutadapt and quality trimmed the 3’ end with a quality threshold of 23 and a minimum length of 35bp using sickle ^49^. We applied the FRAMA pipeline ^50^to assemble the trimmed RNA-seq data to an annotated mRNA assembly, incorporating Trinity ^51^as the assembler. The domestic cat transcriptome served as reference for transcript gene symbol assignment, based on best bidirectional BLASTn hits against CDSs. The resultant orthologous reference transcript with the best aligned hit was used for scaffolding, CDS inference, and splitting of fusions. As output, we retrieved the resultant transcript sequences in the 5’ to 3’ orientation.

### Repeat masking and gene annotation

Next, we carried out repeat and gene annotations on the four genomes. We first masked repeats in each genome using a combination of *ab initio*repeat finding and homology-based repeat annotation using RepeatModeler (http://www.repeatmasker.org) and RepeatMasker (http://www.repeatmasker.org), respectively. For homology-based repeat annotation we used the mammal repeat consensus sequences from Repbase ^52^. During this step we did not mask simple repeats beforehand to improve mapping during the homology-based annotation. We annotated genes using *ab inito*gene prediction, as well as protein and transcriptome homology using the pipeline Maker2 ^53^. Simple repeats were soft-masked using Maker2 ^53^, to allow for more efficient mapping during gene annotation. *Ab initio*gene prediction was carried out using SNAP ^54^and Augustus ^55^. For the protein homology-based annotation step, we combined proteins from the previously annotated domestic cat (*Felis catus*; GCF_000181335.2) and domestic dog (*Canis lupus familiaris*; GCF_000002285.3). We further used a spotted hyena transcriptome for the homology-based annotation step (provided to Maker using the parameter “est=“ for the annotation of the spotted hyena, and “altest=“ for all other hyena species).

### Species tree and molecular dating

For producing a dated phylogenetic tree, we downloaded the assembled transcripts of two additional species from within Feliformia, cheetah (*Acinonyx jubatus*, GCF_003709585.1), and meerkat (*Suricata suricatta*, GCF_006229205.1), and the dingo (*Canis lupus dingo*, GCF_003254725.1) as outgroup. We searched for 1:1 orthologous gene sequences found in all four hyena genomes, as well as the three outgroup species using ProteinOrthov5.11^56^. Preliminary coding sequence alignments were built for each ortholog using MACSE v2.03 ^57^. Fast trees were built using maximum likelihood and the GTR+Γ substitution model in IQTREE v2.0 ^58^. These trees were fed to TreeShrink ^59^for identifying any sequences that could mislead branch length estimates due to difficulties in assembly, annotation, or alignment. Orthologs were aligned again by excluding the flagged taxa. Codons with more than 50% missing taxa were excluded, and all regions were checked by eye for major alignment issues. Possible violations to the assumptions of the GTR+Γ model were tested using PhyloMAd ^60^and IQTREE ^61^, and any loci flagged as model-inadequate were excluded from further analyses. This processing led to 1,219 loci with >1.7M aligned sites.

Using the filtered sequence alignments, we implemented two approaches to estimating the species tree of hyenas. We first estimated phylogenetic trees for each locus using a maximum likelihood and the best-fitting substitution model from the GTR+F+Γ+I family ^62^using IQTREE^63^. Branch supports were calculated using an approximate likelihood ratio test ^64^. These trees were used for subsequent species tree estimation under the multispecies coalescent in ASTRAL v5.6 ^65^, with analyses replicated using either the raw locus trees or by collapsing branches with statistical support <50. We also concatenated all loci to perform a supermatrix approach assuming that gene tree discordance is entirely due to the stochastic inference error associated with having a finite sample size. This analysis was partitioned by locus with independently selected substitution models and assuming proportional variation in branch lengths across loci ^66^.

Molecular dating analyses were performed on the ASTRAL species tree using three fossil calibrations with soft maximum bounds. We constrained the age of the common ancestor of the brown and striped hyena to occur between 4.05 Ma based on the earliest *Parahyaena*fossil (*P. howelli*) ^2,67,68^and 5.2 Ma based on the most recent putative *Hyaena/Parahyaena*ancestor, *Ikelohyaena abronia*^69^. A second constraint was placed on the Hyaenidae-Felidae split to occur between 29 and 35 Ma, based on the reasoning described by Barnett et al. ^70^. Lastly, we constrained the basal divergence of the carnivore clade, assuming it occurred prior to 37.1 Ma based on the age of the fossil *Hesperocyon gregarius*^71^.

Bayesian dating analyses were performed using MCMCtree in PAML v4.8 ^72^, with improved efficiency by implementing approximate likelihood computation ^73^. Loci that were discordant with the species tree topology were excluded from dating analyses to avoid violation of the tree prior^74^and misleading branch length estimates^75^. The remaining loci were partitioned into each of the three codon positions. The molecular evolution of each codon position was described by a GTR+Γ substitution model and an uncorrelated gamma prior on rates across lineages, while a birth-death process was used as the prior for the branching times. We estimated the posterior distribution using Markov chain Monte Carlo (MCMC) sampling. We drew MCMC samples every 1×10^3^steps over 1×10^7^steps, excluding a burn-in phase of 1×10^6^steps. Convergence to the stationary distribution was verified by comparing the parameter estimates of two independent runs. All parameters were found to have effective sample sizes above 1000, as estimated using the R package coda ^76^.

Additional analyses were performed to disentangle any differences between species tree estimation methods (multispecies coalescent versus supermatrix ML method). We estimated the frequency of gene-trees that included each of the tree-quartets observed in the estimated species-trees, also known as gene concordance factors ^77^. Similarly, site concordance factors measure the proportion of sites that support quartets estimated in the species tree ^10^. These two are measures of the decisiveness of the data for a particular phylogenetic resolution, and provide a comprehensive description of disagreement across the data. Concordance factors can also be compared with the support for the two alternative resolutions of each quartet, known as the two discordance factors. We obtained gene- and site concordance factors using IQTREE ^10^from the tree estimated using the multispecies coalescent, after confirming that the concordant and discordant quartets of this estimate also included all the resolutions present in the species tree estimate obtained from concatenation of loci.

In order to more comprehensively explore the history of the hyena lineages, we performed a Bayesian species tree estimation allowing for an event of reticulation under the multispecies network coalescent model ^11,12^. We sampled gene trees and their coalescent times from the full set of locus alignments as implemented using the MCMC_SEQ function of PhyloNet v3.8 ^78^, including exclusively sequences from the four species of hyenas. The posterior distribution was estimated using Markov chain Monte Carlo (MCMC) sampling with samples drawn every 5×10^4^steps across 10^8^steps. A burn-in phase of 2×10^7^steps was excluded, and convergence to the stationary distribution was verified by confirming that all parameters had effective sample sizes above 200, using the R package coda ^76^. We verified that results were identical across two independent runs, and reported the maximum a posteriori (MAP) network from the output.

To further investigate the presence of gene flow between the different lineages we used the D3 statistic ^9^. D3 makes use of pairwise distances and does not require an outgroup genome to polarise between ancestral and derived alleles. D3 uses the topology ((A,B),C) and the equation (BC-AC)/(BC+AC) to uncover differences in branch lengths that may indicate relatively higher levels of gene flow between taxon A and C or taxon B and C. We computed this twice independently with all individuals mapped to either the spotted hyena assembly or striped hyena assembly and only included scaffolds greater than 1Mb in length. We computed the consensus bases pairwise distances −doIBS 2 using non-overlapping sliding windows of 1Mb in length using ANGSDv0.913^13^and specifying the following parameters; output a distance matrix (-makeMatrix 1), −minMapQ 25, −minQ 25, −uniqueOnly 1, −remove_bads 1, only consider sites with at least 5x read depth (-setmindepthind 5), only consider sites found in all four individuals (-minind 4), and only include scaffolds not aligning to the sex chromosomes and larger than 1Mb in length (-rf). We calculated a p-value for each pairwise comparison to evaluate the difference from 0 by calculating the mean, standard deviation, and assuming a normal distribution in R v3.6 ^79^using the pnorm function.

### Sex chromosome alignments

For downstream genetic diversity and demographic history analyses, we first needed to determine which scaffolds were most likely autosomal in origin. To do this we found putative sex chromosome scaffolds for the two *de novo*assembled genomes and removed them from future analyses. We found putative sex chromosome scaffolds by aligning the assemblies to the domestic cat X (Genbank accession: NC_018741.3) and human Y (Genbank accession: NC_000024.10) chromosomes. Alignments were performed using satsuma synteny ^80^and utilising default parameters.

### Ka/Ks calculation

To determine branch specific selection in hyenas we estimated ω-values (Ka/Ks substitution ratios) for 9,400 1:1 orthologs in hyenas (aardwolf, spotted hyena, striped hyena, and brown hyena) and the domestic cat (Ensembl v90 annotation ^81^). We employed a previously described strategy ^82^and performed selection analysis on 1,319 complete genes that did not contain any frameshift indels in the alignment and the longest stretch of at least 200 uninterrupted aligned bases from the other 8,081 genes. Briefly, we used the CODEML program under a branch model ^72^twice independently, once setting the aardwolf as the foreground branch, and the other with all scavengers (spotted, striped, and brown hyenas) as foreground branches. This model was compared via a likelihood ratio test (1 degree of freedom) to the one-ratio model (model=0, NSsites=0) used to estimate the same ω ratio for all branches in the phylogeny. To access functionality of the genes under positive selection, we further looked into the associated MGI phenotypes (Genecards.org) of genes showing highly significant signs of positive selection (X^2^> 7.9 (p=0.005)) in these lineages.

### Olfactory receptor expansions

To estimate the “olfactory ability” of extant Hyaenidae, we curated the olfactory receptor repertoire in the hyena genomes using established methodologies, as earlier described ^83,84^and as performed in a previous study ^82^. To this end, we employed a multi-step process as follows: i) Functional ORs of various mammalian genomes (dog, mouse, opossum, horse, and human) were downloaded from previous studies ^85–88^. These functional ORs were utilized to generate manually curated alignments (of <=40% identity) that served as seeds to build discrete HMM profiles using the HMMER package ^89^. These HMM profiles were utilized for standalone HHM searches against the curated proteome datasets of all Hyaenidae. All recovered hits within the default inclusion threshold (0.01) were considered as putative ORs. ii) Conversely, the proteome datasets of all Hyaenidae were utilized for standalone Pfam search ^90^and RPS-BLAST search as implemented in the CDD database ^91^. All sequences that recovered the 7tm_4 domain (HMM profile for the olfactory domain in the Pfam database, Pfam-ID: PF13853) and cd13954 (PSSM for the olfactory domain in CDD database) as their best scoring hits, in the respective searches, were considered as putative ORs. iii) Putative ORs obtained from steps 1 and 2, and functional ORs downloaded from previous studies were clustered using CD-HIT ^92^into groups at 40% sequence identity threshold and selected candidate sequences from each group were utilized as seeds (diverse starting points) for several standalone TBLASTN searches (cutoff E-value of 1×10^−9^) against individual genomes of all Hyaenidae. These searches recovered additional hits and ensured the recovery of hidden or missed ORs from any unannotated nucleotide sequences in the gene prediction/annotation pipelines. iv) A non-redundant dataset was obtained from steps 1-3 using CD-HIT (97% identify), and these putative ORs were first reviewed using PSI-BLAST searches ^93^against the NCBI-NR database and were further verified through phylogenetic analysis using non-olfactory Class-A GPCRs as outgroup ^83,84^. Protein sequences of all hyenas that recovered 7tm_1 (Pfam-ID: PF00001) as their best hit in the standalone Pfam search were obtained and categorized as Class-A non-olfactory GPCRs and utilized for Maximum-Likelihood (ML) trees computed using the IQ-TREE software ^63^. In all ML trees, the putative ORs formed an unambiguous monophyletic clade with high bootstrap support (>90%), distinct from the non-OR GPCRs.

To assign sub-type (Type-I and Type-II ORs) and sub-group level classification (α, β, γ, etc.) ^85^of all identified ORs from each hyena genome, BLASTP searches were performed against a curated standalone BLAST database comprising previously classified ORs from vertebrate genomes ^85–88^. The database included all identified and classified vertebrate ORs from previous studies, and sequences were tagged based on their category (such as α, β, and γ). Based on the BLASTP searches, all ORs were putatively categorized into each group, and the classification was further substantiated using phylogenetic comparisons.

For comparative analysis of ORs among carnivoran (dog, cat, and tiger) and mammalian genomes, we utilized the previously classified and functional ORs as aforementioned. The cat and the tiger genomes (for which the previous OR annotation is unavailable) were downloaded from Ensembl (*Felis catus*_8.0: GCA_000181335.3, release-92) and NCBI databases (*Panthera tigris*: GCA_000464555.1), respectively, and ORs were identified using the above procedures. To provide a comparative perspective on the OR expansion in the extant Hyaenidae relative to other vertebrates, we constructed an ML tree using full-length intact type-1 ORs (α, β and γ group olfactory receptors) from all analyzed hyenas, carnivoran (dog, cat, and tiger) and other mammalian genomes (human and mouse). The best-fit substitution model for the curated alignment of ORs was estimated using ModelFinder ^62^incorporated in the IQ-TREE software. ML tree topologies were derived using the edge-linked partition model as implemented in the IQ-TREE software, and branch supports were obtained using the ultrafast bootstrap method (1000 replicates) ^94^.

### *De novo*gene repertoire expansion analysis

To cross validate genes under positive selection and the OR repertoire expansions in hyenas, we conducted a secondary *de novo*approach to determine gene families with putative repertoire expansions. This was done to avoid the influence of differing assembly or annotation quality. For this we blasted ORFs found in the raw sequencing reads against the dog proteome (Uniprot ID: UP000002254). Raw forward reads of six carnivores (*Canis lupus*(SRA accession code: SRR8926752), *Panthera tigris*(SRA accession code: SRR5591010), *Ursus arctos*(SRA accession code: SRR830337), *Suricata suricatta*(SRA accession code: SRR11434616), *Paradoxurus hermaphroditus*(SRA accession code: SRR11431891), *Cryptoprocta ferox*(SRA accession code: SRR11097184) and our hyenas were quality checked, converted to fasta, and all ORFs with a minimal length of 90 were extracted and blasted to the dog proteome (Supplementary figure S4). We additionally included hyena genomic data for training the downstream machine learning model. These included *Parahyena brunnea*(concatenation of multiple low coverage genomes: SRA accessions: SRR5886631, SRR5886634, SRR5886635, SRR5886638, SRR5886630, and SRR5886636), *Hyaena hyaena*(SRA accession: SRR11430567), and *Crocuta crocuta*(SRA accessions: Namibia: SRR9914662, Ghana: SRR9914663). ORF extraction was performed using EMBOSS getorf ^95^and protein to protein blasting was done using PLASTp ^96^reporting hits with an e-score over 1×10^5^. The resulting alignment tables were processed in R to remove secondary hits of each read, and normalized by the size of each library. The average size and coverage of the sequencing of the libraries was calculated, using the “estimate size factors” implemented in the DESeq2 R package ^97^, on a set of ∼170 highly conserved BUSCO ^98^genes in the metazoa ortholog set (Supplementary figures S5 and S6). The normalized gene set was analysed with the machine learning randomforest algorithm ^99^(R package randomforest with option ntree=100000 and otherwise default options) to find expanded gene repertoires unique to the Hyaenidae, as well as to the aardwolf, and to the bone-cracking lineage. The 200 genes showing the highest levels of differentiation were further analysed for GO terms.

### Genetic diversity

We used two different methods to estimate the heterozygosity of the four hyena genomes included in our study. To be able to directly compare to previously published heterozygosity results from a wide range of mammalian species ^6,14^, we followed the parameters first set out by Westbury et al, 2018. In brief, this was performed using ANGSD on the mapped bam files subsampled down to 20x (-downsample parameter). We applied the following filters, calculate genotype likelihoods using the SAMtools algorithm (-GL 1), −setMinDepthInd 5, −minmapq 25, −minq 25, −uniqueonly 1, adjust quality scores around indels (-baq 1), produce a folded SFS as output (-fold 1), and remove all scaffolds of less than 1Mb and that aligned to the sex chromosomes (-rf command). We performed this using all species mapped to the striped hyena assembly apart from the spotted hyena which was mapped to the spotted hyena assembly. We further calculated genome-wide heterozygosity and runs of homozygosity using ROHan ^15^. We did this for all species twice independently, once with all genomes mapped to the striped hyena assembly, and once mapped to the spotted hyena assembly. We ran ROHan using default parameters which specifies a window size of 1Mb and ROH if said window has an average heterozygosity of less than 1×10^−5^. We additionally reran the ROHan analysis specifying a ROH cutoff of 5×10^−5^with the aardwolf mapped to both the spotted and striped hyena assemblies, spotted hyena mapped to the spotted hyena assembly, and both the brown and striped hyena mapped to the striped hyena assembly.

### Demographic history

To investigate the respective demographic histories of each species, we performed a pairwise sequentially markovian coalescent model (PSMC)^16^on the diploid genomes of the four hyena species. For this we used all species mapped to the striped hyena assembly apart from the spotted hyena which was mapped to the spotted hyena assembly. We called diploid genome sequences using SAMtools and BCFtools ^100^, specifying a minimum quality score of 20, a minimum coverage of 10, and a maximum coverage of 100. We removed scaffolds found to align to sex chromosomes in the previous step and scaffolds shorter than 1Mb during the diploid sequence construction step. We ran PSMC specifying atomic intervals 4+25*2+4+6 and performed 100 bootstrap replicates to investigate support for the resultant demography. To calibrate the plot we calculated a Hyaenidae average mutation rate. We did this by calculating the average pairwise distance for each species pair and dividing that by 2 × our estimated mean divergence times calculated above (Supplementary tables S9 and S10). We calculated the pairwise distances twice independently, once with all species mapped to the spotted hyena and once with all the species mapped to the striped hyena and took the average of these numbers. We used ANGSD to calculate the pairwise distances using a consensus base call approach (-doIBS 2), and applied the following filters: −makeMatrix 1, −minMapQ 25, −minQ 25, −uniqueOnly 1, −remove_bads 1, −setmindepthind 5, −minind 4, and only include scaffolds not aligning to the sex chromosomes and larger than 1Mb in length. The average Hyaenidae mutation rate was taken from the average of these numbers. This gave us a mutation rate of 7.7×10^−10^per year, only slightly slower than the commonly implemented human mutation rate of 1×10^−9^per year ^16^. To calculate the generational mutation rate we assumed a generation length of six years for all species ^6,7^giving a generational mutation rate of 4.6×10^−9^.

## Acknowledgments

We would like to thank Andrew Heidel, Pablo Santos, Marion East, Heribert Hofer, Oliver Hoener, Bettina Wachter, Simone Sommer, Martin Bens, and Matthias Platzer for providing the assembled spotted hyena transcriptome to aid in annotating the genomes. We also thank Dorina Menegheni and Bernd Senf for the transcriptome data transfer. We thank Armanda Bastos for allowing the use of her laboratory for the aardwolf DNA extractions. We also acknowledge sequencing support from the Swedish National Genomics Infrastructure (NGI) at the Science for Life Laboratory, which is funded by the Swedish Research Council and the Knut and Alice Wallenberg Foundation, and UPPMAX for access to computational infrastructure.

## Author Contributions

Conceptualization, MVW, MH; Formal analysis, MVW, DLD, DAD, AK, SP, JHG; Investigation, MVW; Writing – Original Draft MVW; Writing – Review & Editing MVW, LW, FD, MH; Resources, KN, FD, LD, MH; Data curation; MVW, SR, AW; Supervision, TS, MH; Funding Acquisition, MH;

## Data availability

An NCBI SRA accession code for the raw aardwolf data will be available upon acceptance.

