## Supplementary information for "Ecological specialisation and evolutionary reticulation in extant Hyaenidae"

**Supplementary figures**

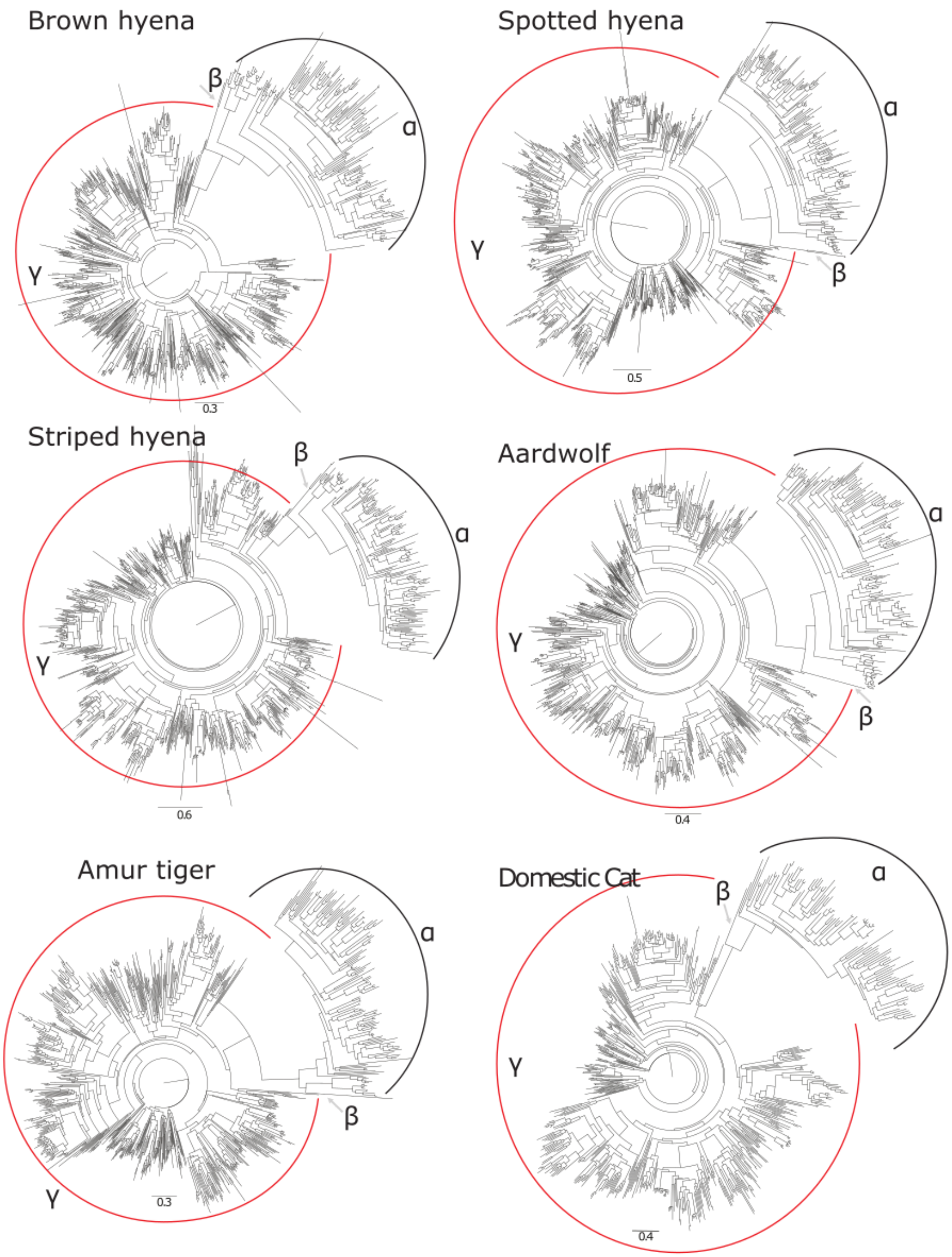


**Supplementary Figure S1:** Maximum likelihood (ML) tree topologies showing the relationships between the different subfamilies of olfactory receptor (OR) repertoires from each species. Tree topologies are consistent with earlier studies separating the λ and α subfamily ORs, while the β subfamily ORs clustered basal to the clade clustering the α. The scale bar represents the number of amino-acid substitutions per site.

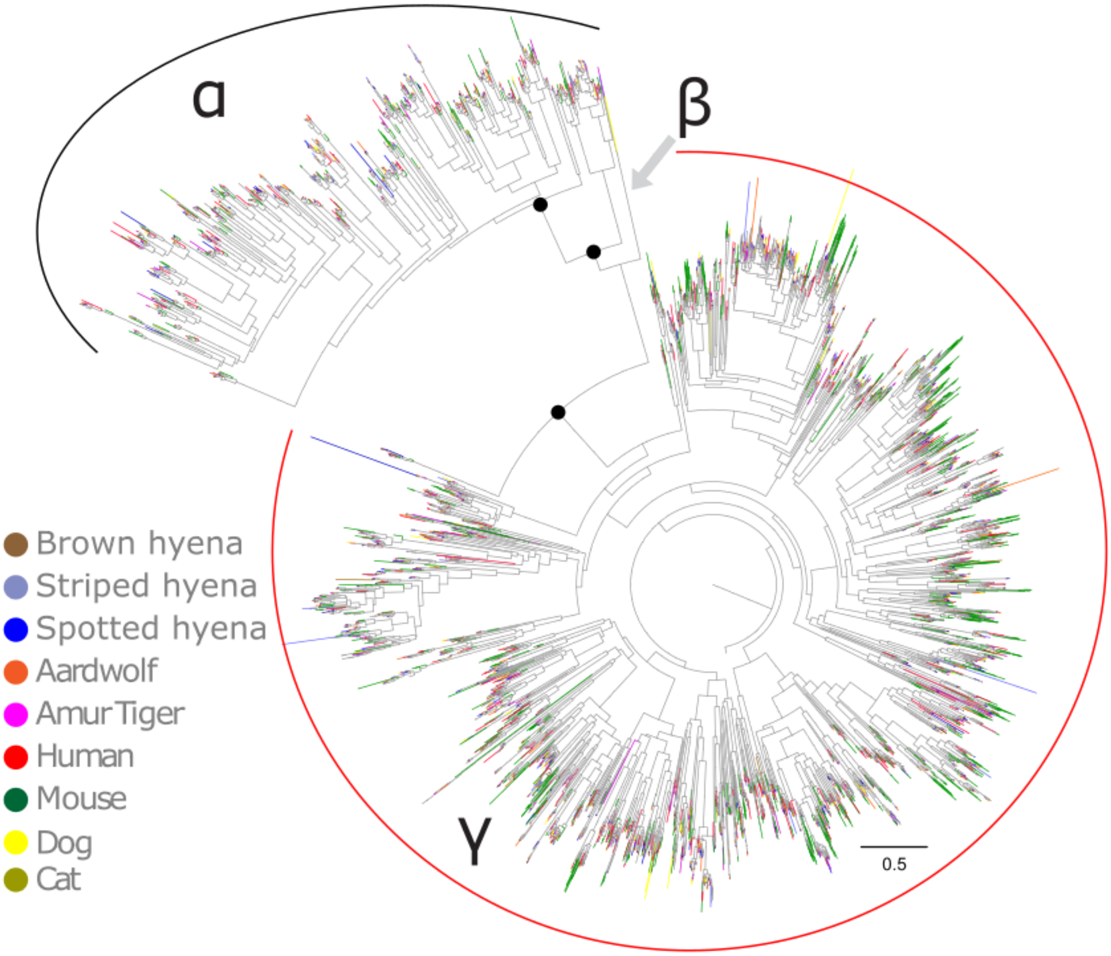


**Supplementary Figure S2:** The inferred Maximum likelihood (ML) tree topology for a comparative perspective on the OR expansion in the extant Hyaenidae relative to other vertebrates. ML tree was constructed using the edge-linked partition model as implemented in IQ-TREE employing full-length and intact ORs from all analyzed hyenas, carnivoran (dog, cat, and tiger) and other mammalian genomes (human and mouse). The best-fit substitution model for the curated alignment of ORs was estimated using ModelFinder and JTT+F+R10 was chosen as the best-fit according to Bayesian Information Criterion (BIC) as implemented in IQTREE. Branch supports were obtained using the ultrafast bootstrap method (1000 replicates) and the red dot indicates confidence estimates (95 % bootstrap support) for the nodes that distinguish α and γ ORs. The scale bar represents the number of amino-acid substitutions per site.

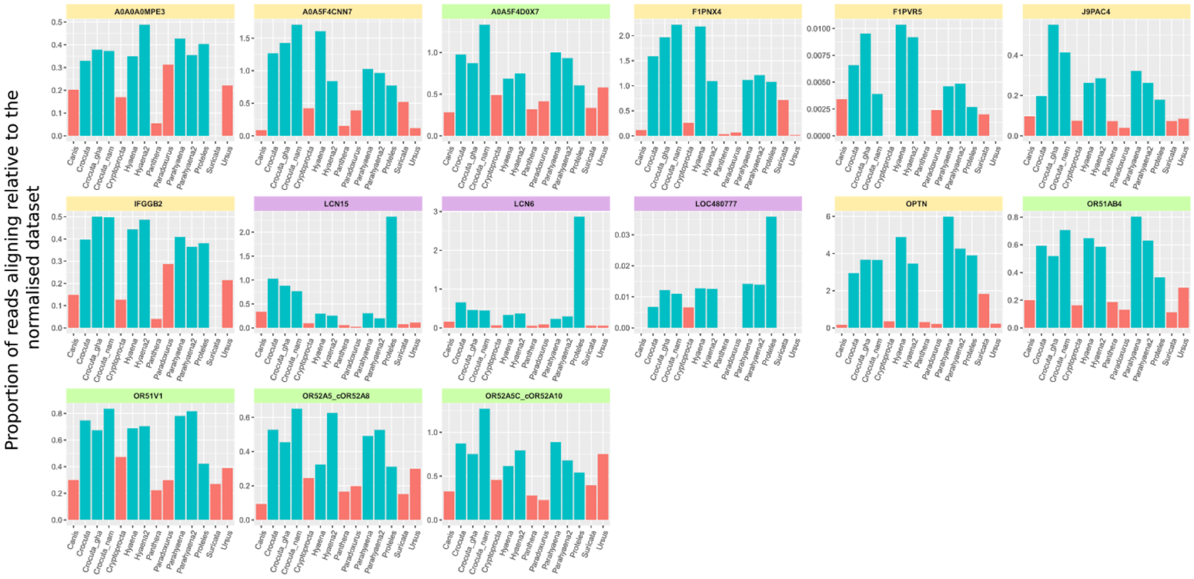


**Supplementary Figure S3:** Genes with putative repertoire expansions as identified by randomforest with known functions in immune response (yellow header), olfactory (green header) and responses to toxins (purple header).

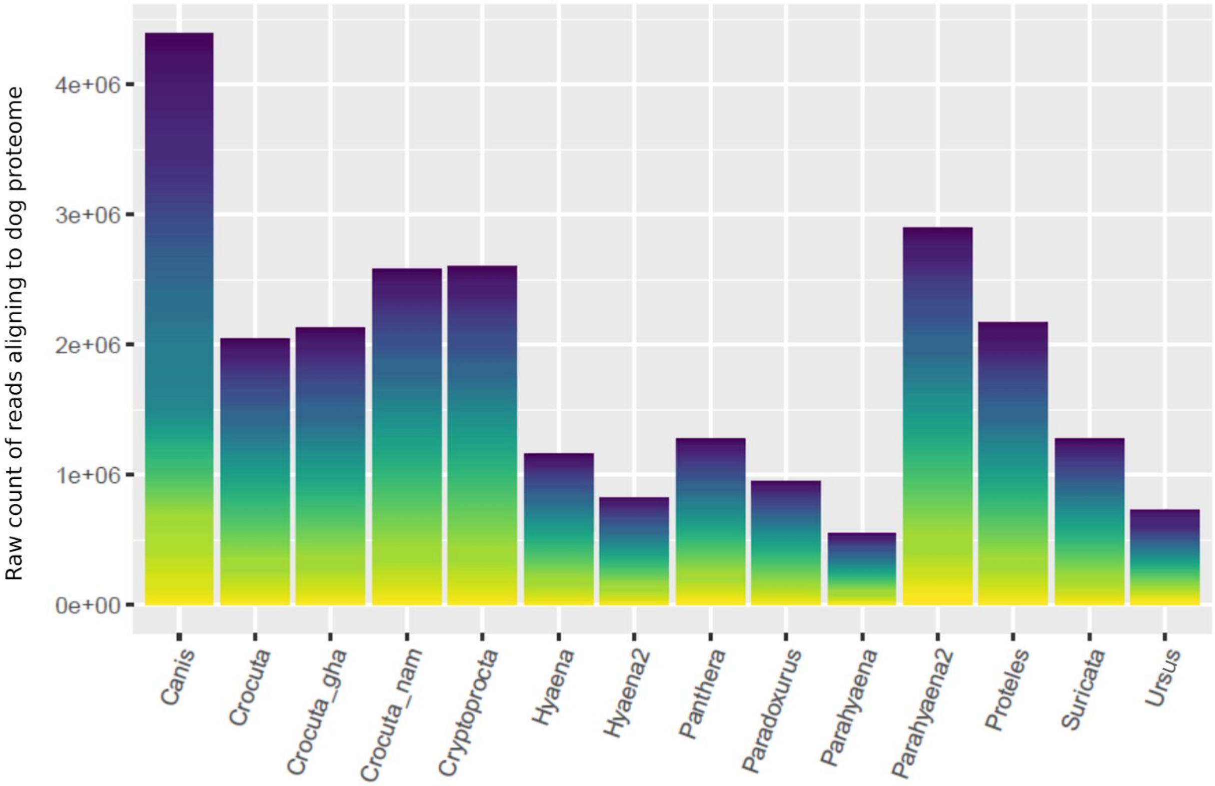


**Supplementary Figure S4:** Sum of raw counts for the number of reads aligning to all genes in the dog proteome from each library.

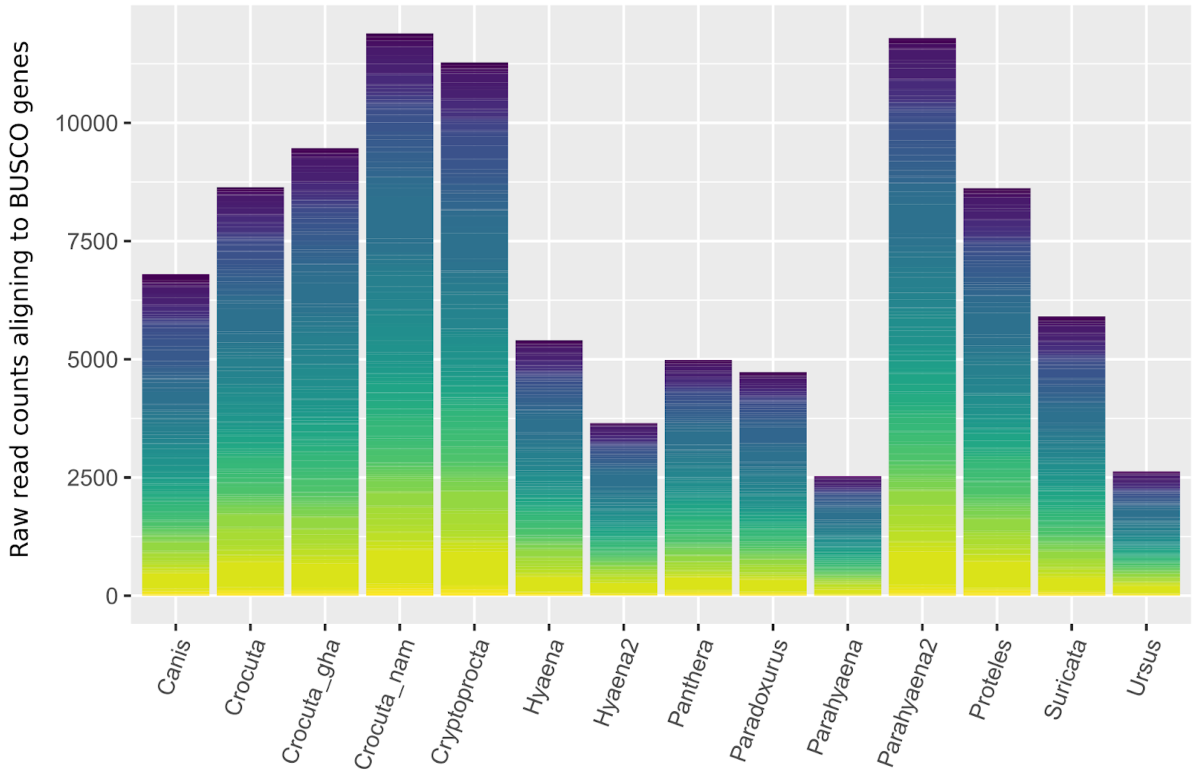


**Supplementary Figure S5:** Sum of raw read counts aligning to all selected metazoan BUSCO genes in each library.

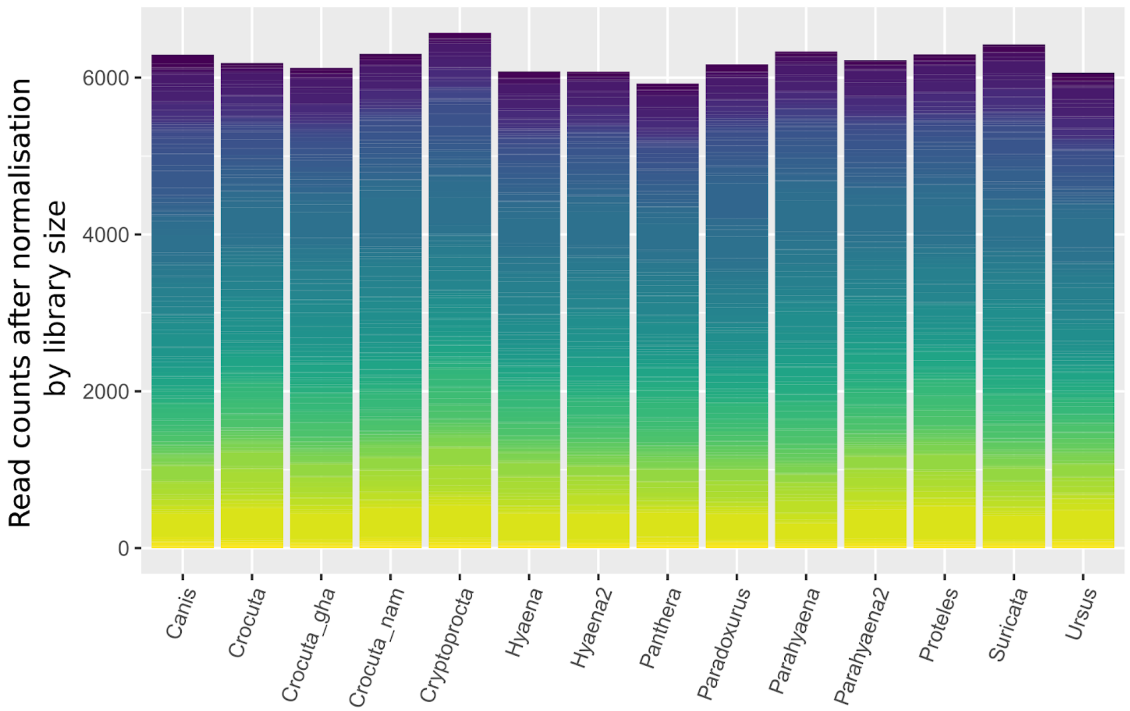


**Supplementary Figure S6:** Sum of library size after normalisation with the metazoan BUSCO gene database.

**Supplementary tables**

**Supplementary table S1:** D3 results mapping to the aardwolf.

| Topology tested | Mean D3 | St. deviation | p-value |
|---|---|---|---|
| ((Spotted,Brown),Aardwolf) | -0.01360466 | 0.006422542 | 0.03415242 |
| ((Striped,Brown),Aardwolf) | 0.00228674 | 0.002343745 | 0.3292242 |
| ((Striped,Spotted),Aardwolf) | 0.01589084 | 0.006459946 | 0.01389746 |
| ((Striped,Brown),Spotted) | -0.003162934 | 0.003845739 | 0.4108199 |

**Supplementary table S2:** D3 results mapping to the spotted hyena

| Topology tested | Mean D3 | St. deviation | p-value |
|---|---|---|---|
| ((Spotted,Brown),Aardwolf) | -0.02185175 | 0.005818448 | 0.0001729289 |
| ((Striped,Brown),Aardwolf) | 7.22E-05 | 0.00228972 | 0.9748423 |
| ((Striped,Spotted),Aardwolf) | 0.0219239 | 0.00593843 | 0.0002226133 |
| ((Striped,Brown),Spotted) | -5.67E-05 | 0.003848203 | 0.9882512 |

**Supplementary table S3:** Genes under positive selection in the bone-cracking hyena lineage

- Attached as a spreadsheet

**Supplementary table S4:** Genes under positive selection in the aardwolf lineage

- Attached as a spreadsheet

**Supplementary table S5:** Autosomal wide average levels of heterozygosity following the parameters set out in Westbury et al 2018, 2019. Numbers from all non-hyena species were taken from Westbury et al 2018, 2019.

| **Species** | **Autosome-wide heterozygosity** |
|---|---:|
| Yellow baboon | 0.001680 |
| Chimpanzee | 0.001080 |
| Human (African) | 0.000791 |
| Aardwolf | 0.000663 |
| Human (European) | 0.000595 |
| Spotted hyena | 0.000591 |
| Panda | 0.000497 |
| Bonobo | 0.000468 |
| Bowhead whale | 0.000357 |
| Walrus | 0.000354 |
| Polar bear | 0.000301 |
| Beluga | 0.000291 |
| Cheetah | 0.000269 |
| Orca | 0.000214 |
| Iberian Lynx | 0.000176 |
| Narwhal | 0.000142 |
| Island fox (San Miguel) | 0.000139 |
| Striped hyena | 0.000135 |
| Brown hyena | 0.000112 |

**Supplementary table S6:** ROHan results mapping to the striped hyena and implementing default parameters

| | Aardwolf | Brown hyena | Spotted hyena | Striped hyena |
|---|---|---|---|---|
| **Segments in ROH(%)** | 0 | 0 | 0 | 0.050917 |
| **Avg. length of ROH (bp)** | 0 | 0 | 0 | 1000000 |
| **Global heterozygosity rate:** | 0.002399 | 0.000555 | 0.001511 | 0.000753 |
| **Lower 5%** | 0.002258 | 0.000485 | 0.001350 | 0.000671 |
| **Upper 5%** | 0.002556 | 0.000628 | 0.001643 | 0.000839 |

**Supplementary table S7:** ROHan results mapping to the spotted hyena and implementing default parameters

| | Aardwolf | Brown hyena | Spotted hyena | Striped hyena |
|---|---|---|---|---|
| **Segments in ROH(%)** | 0 | 0 | 1.965160 | 0 |
| **Avg. length of ROH (bp)** | 0 | 0 | 2750000 | 0 |
| **Global heterozygosity rate:** | 0.002338 | 0.000611 | 0.001339 | 0.000939 |
| **Lower 5%** | 0.002206 | 0.000543 | 0.001255 | 0.000847 |
| **Upper 5%** | 0.002498 | 0.000681 | 0.001448 | 0.001027 |

**Supplementary table S8:** ROHan results when specifying a ROH as a 1Mb window with heterozygosity of less than 5e-5 and mapping the aardwolf to both the striped and spotted hyena assemblies, the brown and striped hyena to the striped hyena assembly, and the spotted hyena to the spotted hyena assembly.

| | **Aardwolf (spotted map)** | **Aardwolf (striped map)** | **Brown hyena** | **Spotted hyena** | **Striped hyena** |
|---|---|---|---|---|---|
| **Segments in ROH(%)** | 0 | 0.051047 | 0.357690 | 5.124780 | 2.403340 |
| **Avg. length of ROH (bp)** | 0 | 1000000 | 2333330 | 2017540 | 1840000 |
| **Global heterozygosity rate:** | 0.002339 | 0.002363 | 0.000580 | 0.001380 | 0.000771 |
| **Lower 5%** | 0.002202 | 0.002220 | 0.000516 | 0.001262 | 0.000690 |
| **Upper 5%** | 0.002487 | 0.002527 | 0.000650 | 0.001489 | 0.000856 |

**Supplementary table S9:** Pairwise distances when mapping to the spotted hyena

| | **Aardwolf** | **Brown hyena** | **Spotted hyena** | **Striped hyena** |
|---|---|---|---|---|
| **Aardwolf** | | 0.01861794 | 0.01783186 | 0.01862187 |
| **Brown hyena** | 0.01861794 | | 0.01150387 | 0.004658385 |
| **Spotted hyena** | 0.01783186 | 0.01150387 | | 0.01150623 |
| **Striped hyena** | 0.01862187 | 0.004658385 | 0.01150623 | |

**Supplementary table S10:** Pairwise distances when mapping to the aardwolf

| | **Aardwolf** | **Brown hyena** | **Spotted hyena** | **Striped hyena** |
|---|---|---|---|---|
| **Aardwolf** | | 0.01867753 | 0.01819161 | 0.01876537 |
| **Brown hyena** | 0.01867753 | | 0.01168854 | 0.00468913 |
| **Spotted hyena** | 0.01819161 | 0.01168854 | | 0.01176519 |
| **Striped hyena** | 0.01876537 | 0.00468913 | 0.01176519 | |
